# Identification and characterization of cherry (*Cerasus pseudocerasus* G. Don) genes in response to parthenocarpy induced by GA3 through transcriptome analysis

**DOI:** 10.1101/351775

**Authors:** Binbin Wen, Wenliang Song, Mingyue Sun, Min Chen, Qin Mu, Xinhao Zhang, Xiude Chen, Dongsheng Gao, Wei Xiao

## Abstract

Fruit set after successful pollination is a key process in the production of sweet cherries, but low fruit rate is the main problem for crop production in sweet cherries. Gibberellin treatment can directly induce parthenogenesis without pollination; therefore, gibberellin treatment is a very important method to improve the fruit setting rate of sweet cherries. Exogenous gibberellin can satisfy the hormone requirement during fruit growth and development. Some related studies have mainly focused on physiological aspects, such as breeding, branching, fertilization, etc., and the molecular mechanism is not clear. In this study, we analyzed the transcriptome of ‘Meizao’ sweet cherry fruit treated with gibberellin during the anthesis and hard-core period to determine the genes associated with parthenocarpic fruit set. A total of 765 and 186 differentially expressed genes (DEGs) were found at anthesis and the hard-core stage after gibberellin 3 (GA3) treatment, respectively. The differentially expressed genes between the control and GA3 treatment showed that the GA3 response mainly included parthenocarpic fruit set and cell division. Exogenous gibberellin stimulated sweet cherry parthenocarpy and enlargement, which were verified by qRT-PCR results of related genes and the parthenocarpic fruit set and fruit size. Based on our research and previous studies on *Arabidopsis thaliana*, we identified the key genes associated with parthenocarpic fruit set and cell division. Briefly, we found patterns in the sweet cherry fruit setting-related DEGs, especially those associated with hormone balance, cytoskeleton formation and cell wall modification. Overall, the result provides a possible molecular mechanism regulating parthenocarpic fruit set, which is important for basic research and industrial development of sweet cherries.

**Highlight:** cherry genes in response to parthenocarpy and promote to fruit setting induced by GA3.

## Introduction

Fruit set is an important step in the process of fruit growth and development. Fertilization is a very important step in the fruit setting process, in which the ovary becomes enlarged with the development of the embryo, thus inducing fruit formation [1]. A variety of hormonal synergies play a major role during the control of the fruit set process. When GA, auxin, and cytokinin alone play a role, the fruit can grow and develop to a certain stage, but their synergistic effect can make the fruit develop normally [2,3].

Exogenous spraying of 5 mg / L GA3 can maintain the activity of citrus parietal cells and promote their division, thereby increasing the fruit set rate [4]. Before full bloom, the treatment of grape flower spikes with 30 mg/L GA3 can significantly improve the fruit set rate [5]. When *SIARF7*, a negative regulator of fruit set, was silenced in transgenic tomato plants and *SIGH3* was upregulated, the IAA content decreased in the transgenic plants, the number of middle and endocortical cells decreased, and their volume increased [6,7]. In addition to auxin and gibberellin, the fruit setting rate is also affected by cytokinin, which affects fruit growth by promoting cell division [8]. Gibberellin transcriptome analysis between normally pollinated and parthenocarpic cucumbers showed that auxin, gibberellin and cytokinin interacted in parthenocarpic cucumbers [9]. Therefore, fruit can be formed without pollination, which is parthenogenesis without seeds.

Parthenocarpy is an agronomic trait in which fruit develops without pollination and fertilization under conditions of natural or human intervention [10]. When the external environment is not conducive to pollination and pollen quantity is insufficient or pollen is of poor quality, parthenocarpy can solve the problem of low fruit set. For example, cucumber [11], lemon [4], tomato [12], banana [13], etc. can produce fruit under natural conditions.

During production, the exogenous application of gibberellin can induce parthenocarpy. Adding 1500 mg / L GA3 during anthesis in *Annona lucidum* can significantly increase fruit size and allow high-quality parthenogenetic fruit to be obtained [14]. The addition of 500 mg / L GA_4 + 7_ can not only induce the parthenocarpy of sand pear but also significantly enhance the sugar and sugar ratio [15]. Gibberellins play a major role in inducing parthenogenesis and fruit development in tomato [16]. The overexpression of the citrus gene *GA20ox1* changed the path of gibberellin biosynthesis in tomato, resulting in a high concentration of GA4 and causing the tomato to be uniparental [17,18]. *DELLA* protein represses the transduction of gibberellin signaling. After gibberellins combine with the gibberellin receptor *GID1*, ubiquitination of the *DELLA* protein can be stimulated and can result in 26S proteasome-mediated ubiquitin degradation. Gibberellin synthesis pathway-related gene expression changes in *SIDELLA* silenced transgenic tomato and by promoting ovarian cell division and growth, induced parthenocarpy [19,2].

Sweet cherry, as a fresh fruit, is delicious and juicy. It has become a major industry providing farmers with increased incomes. However, the low fruit setting rate caused by self-fruiting or a low self-setting rate has become a bottleneck restricting the development of this industry. In addition, exogenous gibberellin can induce parthenocarpy in fruit, thereby increasing the fruit setting rate. To describe the molecular mechanism of gibberellin in the process of inducing parthenocarpic fruit set, with the help of third-generation sequencing technology, transcriptome sequencing (RNA-seq) was carried out at two key developmental periods 7 days after GA3 treatment or no treatment in ‘Meizao’ European sweet cherry grown under greenhouse conditions. For the first time, RNA-seq was used to analyze the developmental metabolic pathways and gene expression patterns in sweet cherry fruit after GA treatment. The genes that promote fruit setting, parthenogenesis, cell division and growth are mainly analyzed. The aim of this study is to provide a scientific and theoretical basis for the molecular mechanism of fruit enlargement and fruit setting and to examine the effect of plant hormone regulation on the growth and development of fruit in parthenogenetic European sweet cherry treated by exogenous GA3. Transcriptome sequencing and differentially expressed genes were used to provide a molecular foundation for the induction of parthenocarpic fruit set in sweet cherry by exogenous gibberellin.

## Materials and methods

### Plant material

Trees of the ‘Meizao’ variety of sweet cherry (*Cerasus pseudocerasus* G. Don Meizao) in the anthesis and hard-core stage were treated on February 26 and March 17 at the commercial greenhouse in Jueyu (Tai’an, China). The experimental materials were 10-year-old adult trees of similar condition. All samples were collected from different locations on the same tree. Sweet cherry trees at anthesis and the hard-core stage were sprayed with 300 mg/L of GA3 for the treatment and left under natural conditions for the control. After treatment with gibberellin, 30 cm*25 cm transparent paper bags were used to cover flowers to avoid interference from other pollen and to allow normal management.

The GA3 treatment was performed at day 0 (anthesis with GA treatment, T1, and no GA treatment, CK1), day 10 (the first enlargement period), day 20 (the hard-core period with GA treatment, T2, and no GA treatment, CK2) and day 30 (the second enlargement period), and samples were taken 7 d after each treatment, immediately put into liquid nitrogen for quick freezing, placed in a −80 °C cryogenic refrigerator and kept for future use. In each period, 10 sweet cherries were randomly selected for the determination of transverse diameter and parthenogenetic rate. In this way, two key stages of the development of sweet cherries were examined.

### RNA extraction

The two key developmental periods of sweet cherry were considered. In each critical period, RNA was extracted from sweet cherries that were treated or untreated. Total RNA was extracted using a TIANGEN kit. The quality and quantity of RNA were determined using a Kaiao K5500 spectrophotometer (Kaiao, Beijing) followed by 1% agarose gel electrophoresis (18S and 28S bands). The integrity and concentration of the RNA samples were detected using the Agilent 2100 RNA Nano 6000 Assay Kit (Agilent Technologies, CA, USA), and high-quality RNA was sent to Beijing Genomics for further quality assessment.

### Library preparation for transcriptome analysis

After testing the total RNA samples, the mRNA was enriched with magnetic beads containing oligo(dT) primers. Fragmentation buffer was added to the obtained mRNA to produce short fragments, and then the fragmented mRNAs were used as templates. The first cDNA chain was synthesized using six-base random primers; buffer, dNTPs, RNaseH and DNA polymerase I were added to synthesize the second CDNA chain; and the chain was purified by a QIAquick PCR kit and eluted with EB buffer. The purified double-stranded cDNA was subjected to end repair, base A addition and sequencing, and fragment size was then determined by agarose gel electrophoresis and PCR amplification, thereby completing final sequencing library preparation. The constructed library was sequenced with the Illumina platform.

### Illumina read processing and de novo assembly

Illumina high-throughput sequencing results were originally presented as raw image data files. After CASSAVABase base recognition (base calling), the data were transformed into the original sequence (sequenced reads), and the results were stored in the FASTQ (fq) file format. In FASTQ format, the ASCII code value corresponding to each base quality character minus 33 (Sanger quality system) is the sequencing quality score of the base. Different scores represent different nucleotide sequencing error rates, such as score values of 20 and 30 indicating base sequence error rates of 1% and 0.1%, respectively. To ensure the quality of the data for information analysis, we filtered out the sequences of sequencing linkers, low-quality sequences and reads with an N ratio greater than 5%. After filtering, the total number of bases with a mass value greater than 30 (error rate less than 0.1%) was divided by the total number of bases to obtain the Q30 value; the higher this proportion is, the higher the quality of the sequencing is. Once high-quality clean reads were obtained, a follow-up analysis based on the clean reads was performed.

### Functional annotation and classification

High-quality clean reads were obtained by removing the sequences of sequencing joints, low-quality sequences (Q is less than 19 bases), and reads with an N ratio of more than 5%. Then, TopHat software [20] and Bowtie 2 [21] were used to compare the sequences to the European sweet cherry genome located at https://www.rosaceae.org/species/prunus_avium/genome_v1.0.a1. The number of base mismatches was not more than two, and the annotation results of the genes were obtained by comparing them with the European sweet cherry genome data. In addition, NCBI, UniProt, GO and KEGG were used to annotate the different genes. For GO analysis, the NCBI NR database was first used for BLAST alignment. Blast2GO was used to obtain GO entries corresponding to each gene. Enrichment analysis was performed on each pathway in KEGG to determine the significance of differentially expressed genes (enrichment of pathway). For GO analysis, we first used the NCBI NR database to perform a BLAST comparison. Then, we used Blast2GO to get the GO terms corresponding to each gene. Enrichment analysis was performed on each pathway in KEGG to determine the significance of differentially expressed genes.

### DEG analysis

Sequences were aligned by using TopHat software [20] and Bowtie 2 [21]. The clean reads from the two different developmental stages were obtained through transcriptome sequencing as described above. The complete dataset excluded sequences with more than two base mismatches [22], and the amount of mRNA produced by each transcribed gene was calculated using FPKM [23] to estimate the value of gene expression. The gene expression in samples from the two developmental stages were compared. We used DESeq for differential gene expression analysis and compared the treatment group and reference group. The genes with |log2Ratio| ≥ 1 and q<0.05 were selected as significant differentially expressed genes [24]. The differentially expressed genes were subjected to GO analysis and KEGG metabolic pathway enrichment analysis based on the calculation of the number of genes for each item and results from a hypergeometric test after p-value correction with a threshold of q<0.05. The GO items satisfying this condition are defined those that are significantly enriched in the differentially expressed genes. A pathway defined by a Q-value ≤ 0.05 (corrected p-value) is a significantly enriched KEGG pathway for differentially expressed genes. The definition of a GO item meeting the condition for significant enrichment in differently expressed genes is one having a Q-value ≤ 0.05 (pathway corrected p-value) based on the difference in gene expression enrichment in a KEGG pathway.

### Quantitative real-time PCR (qRT-PCR) validation

To verify the results of the transcriptome analysis, we selected the genes involved in seed development, cell wall development and GA metabolism or signaling. Total RNA was reverse transcribed (1 μg per sample) in a 20 μl reaction using a cDNA reverse transcription synthesis kit (RR047A, TaKaRa, China). The specific primers used for the tested genes are listed in Table S1. The SYBR^®^ Premix Ex Taq TM (Tli RNaseH Plus) kit (Thermo Fisher) was used for the fluorescence quantitative PCR reaction. The reaction system included the following: 12.5 μL SYBR^®^ Premix Ex Taq (2 ×), 1 μL each of forward and reverse primers, 1 μL cDNA, and addition of ddH_2_O to 25 μL. The experimental design included 3 technical replicates. The fluorescence quantitative PCR reaction conditions were as follows: 35 ~ 40 cycles of pre-denaturation at 95 °C for 30 s, denaturation at 95 °C for 5 s, and annealing at 60 °C for 30 s. After the reaction, a fluorescence curve and a melting curve were obtained. Finally, the comparative Ct (2-ΔΔCt) method was used for data analysis [25].

## Results

### Growth and development characteristics of fruit after exogenous GA3 treatment

The full growth and development of fruit was observed after treatment with 300 mg/L GA3 of ‘Meizao’ sweet cherry trees in the anthesis period and the hard-core period. The size of sweet cherry with GA3–treated shown in table 1. After 7 days, the vertical and transverse diameters of trees in the GA3 treatment were significantly higher than in the control. Among them, the increase in the vertical diameter was higher than in the transverse diameter. In the anthesis stage, compared with the control group, the diameter of GA3-treated fruits increased by 123.9%, and the vertical diameter increased by 214.09% (Figure 1). In the first expansion period, the growth rate of the transverse diameter was not significantly different from that in the control, but the vertical diameter reached 34.16%. After the hard nucleus stage, all the fruit of the control group fell off.

**Table.1.**
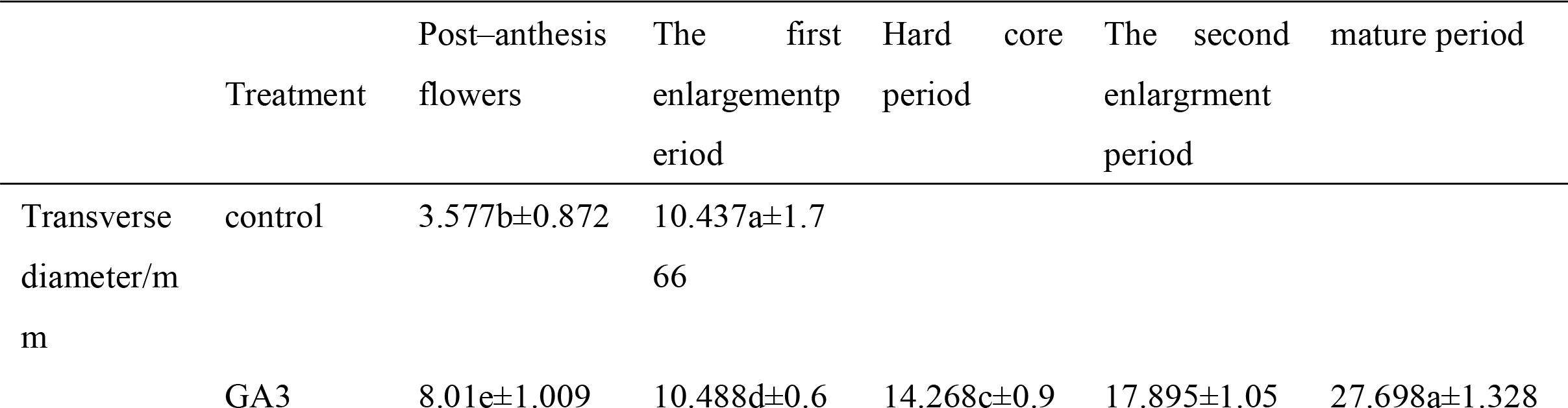
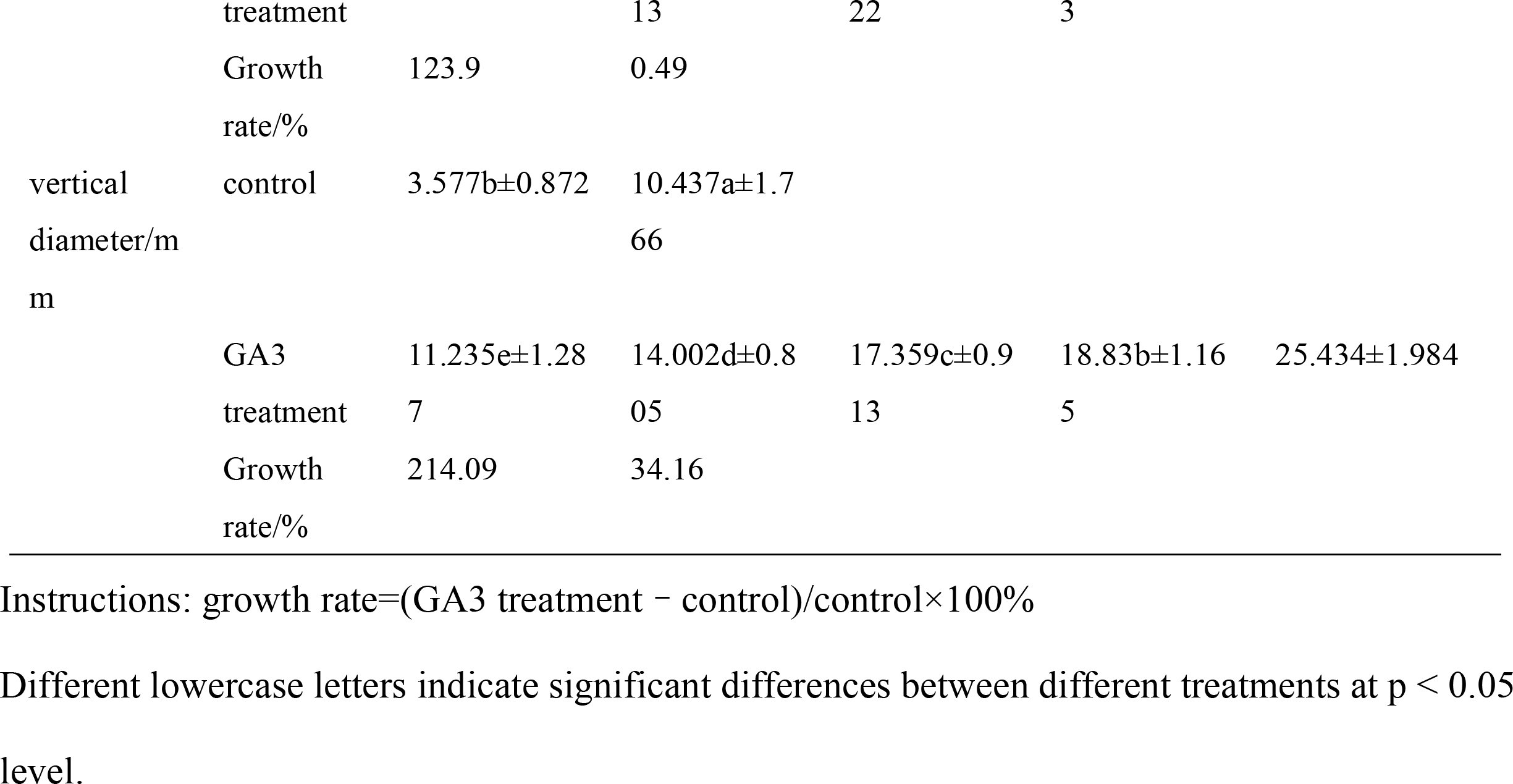
The size of sweet cherry with GA3-treated

**Fig. 1.**
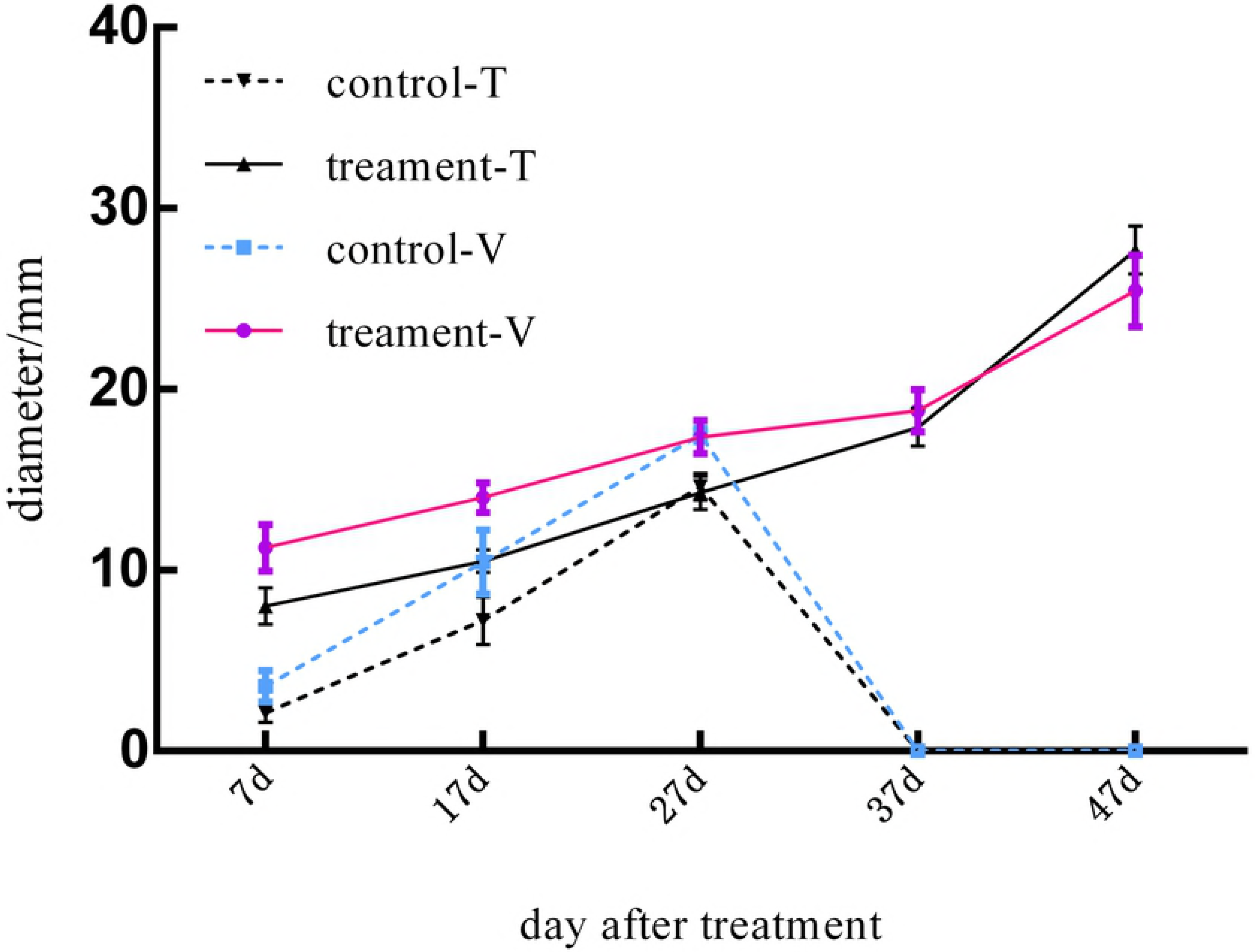
growth curves of 300mg/L-treated and control sweet cherry T, transverse diameter; V, vertical diameter

### Fruit setting rate and parthenogenetic rate of fruit after exogenous GA3 treatment

In the four stages of the development of sweet cherry, 300 mg/L GA3 was applied, and the trees in the natural conditions (control) were not treated. The fruit set rate was 77.33% after a week of spraying GA3 at the anthesis stage, and in natural conditions, it was 12.55%. During the hard nucleus period, the untreated sweet cherries had completely fallen off, while the GA3-treated sweet cherries maintained a fruit setting rate of 57.23%. It can be seen from the figure 2A that spraying GA3 in the falling flower period can both induce parthenogenesis and significantly increase the fruit setting rate. After GA3 treatment, the rate at which fruit dropped from first expansion to the second enlargement was 34.3%, while the parthenogenetic rate was 100% (Figure 2B). With the growth of the fruit, the development of the seeds was inhibited, and the phenomenon of abortion occurred.

**Fig. 2.**
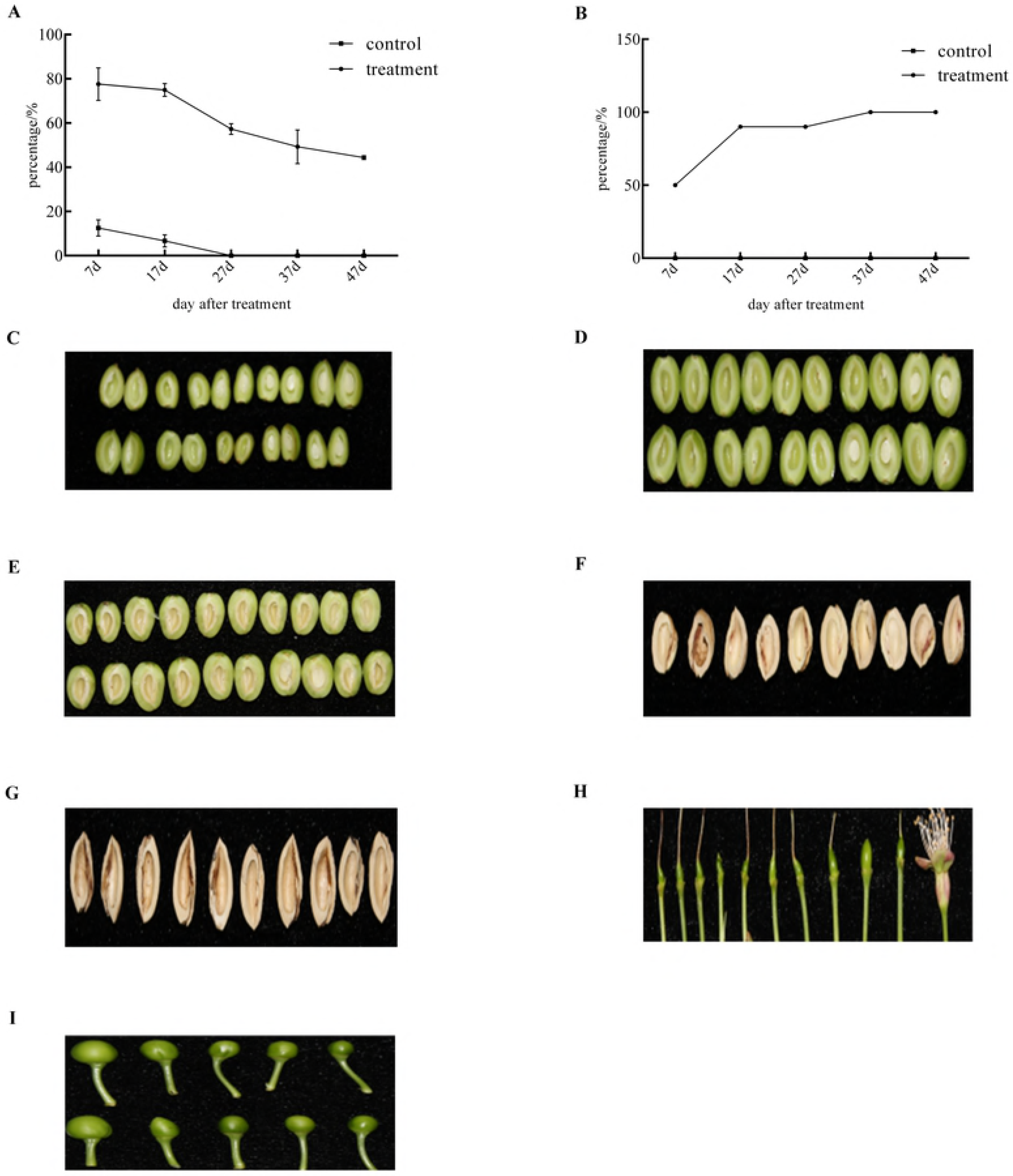
Changes in shape and size of control and GA3 treatment. fruiting rate (A). Parthenogenetic rate (B). Treatment:7days after treatment (DAT) (C), 17 DAT (D), 27 DAT (E), 37 DAT (F), 47 DAT (G). Control:7 days after full bloom (H), 17days after full bloom (I)

### Analysis of transcriptional sequence data

Through Illumina platform sequencing, a total of 137658266, 134689144, 140680232 and 138579108 raw reads were obtained from control and GA-treatment groups, respectively (Table 2). In total, 136440870, 133557948, 139421700 and 137336514 clean reads were selected for further analysis by removing the contaminated reads from the linkers, low-quality reads and reads with an N ratio greater than 5%, The proportion of clean reads to total reads obtained from the four libraries was 99.11%, 99.16%, 99.11% and 99.10%. Among these reads, 83%, 83.67%, 83%, and 79.67% were mapped to the European sweet cherry genome (https://www.rosaceae.org/species/prunus_avium/genome_v1.0.a1), including 81.65%, 80.87%, 80.86%, and 81.22% mapped to the exonic regions; 8.62%, 9.45%, 9.29%, and 9.12% mapped to the intronic regions; and 9.74%, 9.69%, 9.85%, and 9.66% mapped to the intergenic regions.

**Table 2.**
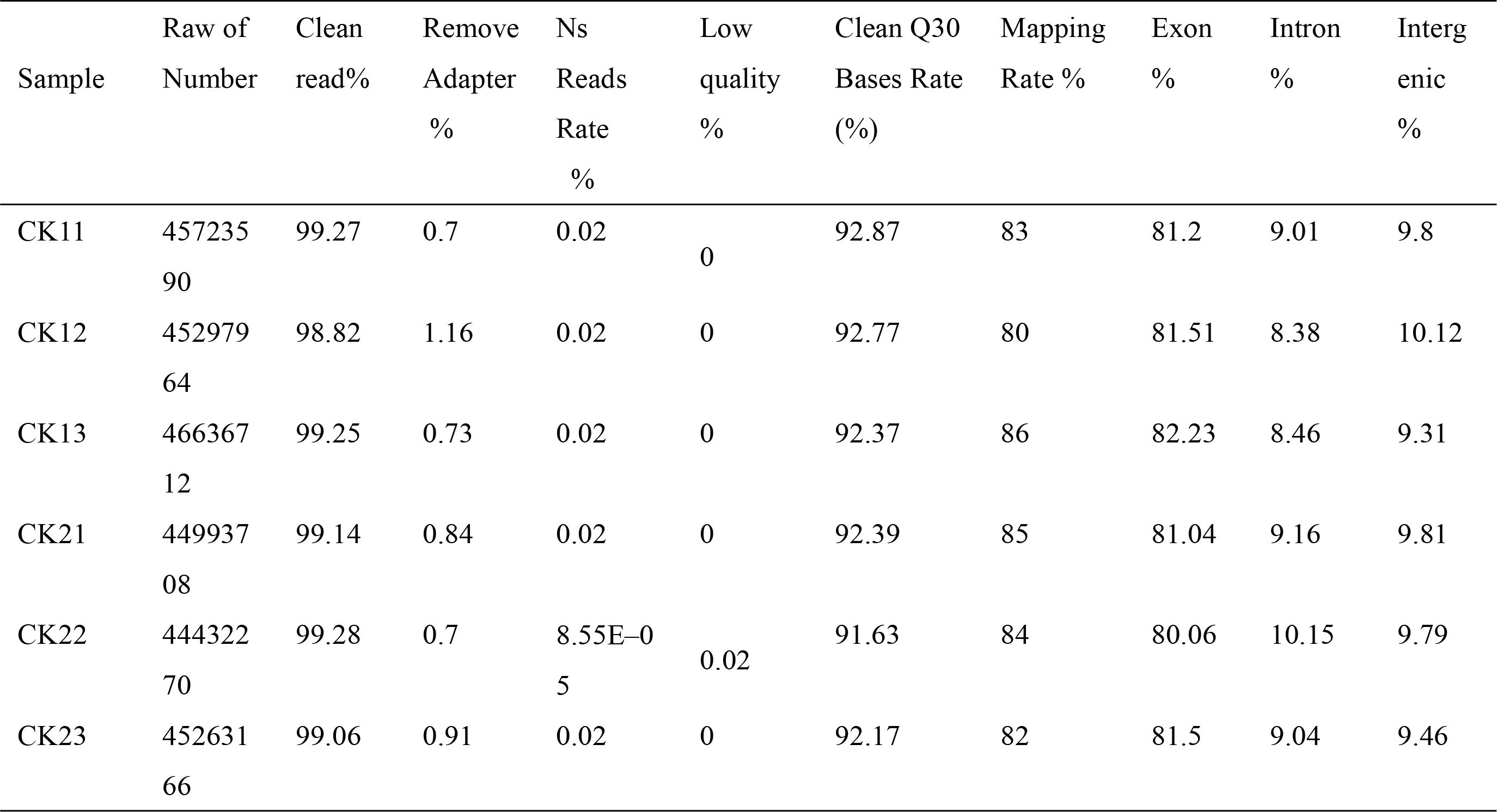
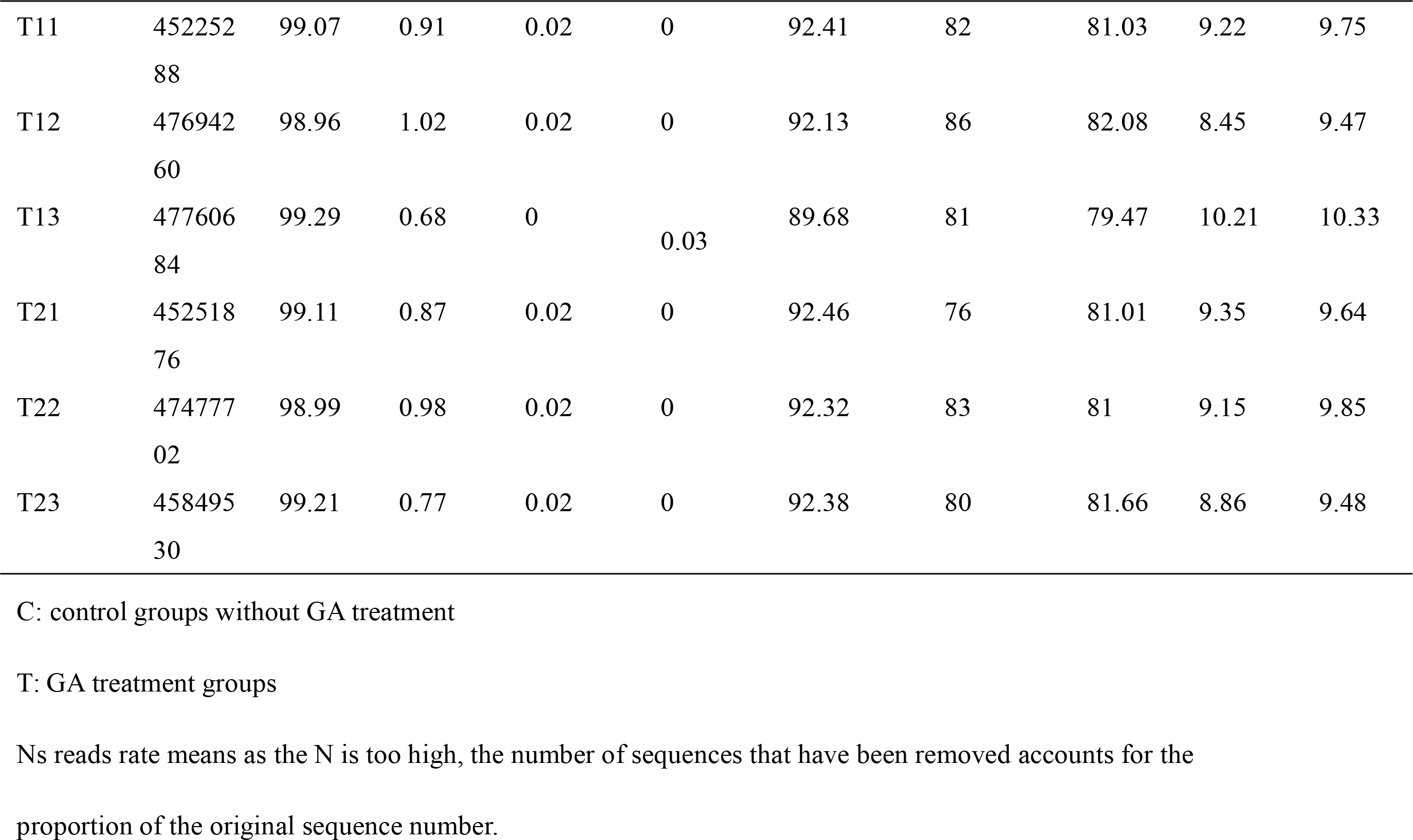
The quality control of the clean data

| Sample | Raw of Number | Clean read% | Remove Adapter % | Ns Reads Rate % | Low quality % | Clean Q30 Bases Rate (%) | Mapping Rate % | Exon % | Intron % | Intergenic % |
| --- | --- | --- | --- | --- | --- | --- | --- | --- | --- | --- |
| CK11 | 45723590 | 99.27 | 0.7 | 0.02 | 0 | 92.87 | 83 | 81.2 | 9.01 | 9.8 |
| CK12 | 45297964 | 98.82 | 1.16 | 0.02 | 0 | 92.77 | 80 | 81.51 | 8.38 | 10.12 |
| CK13 | 46636712 | 99.25 | 0.73 | 0.02 | 0 | 92.37 | 86 | 82.23 | 8.46 | 9.31 |
| CK21 | 44993708 | 99.14 | 0.84 | 0.02 | 0 | 92.39 | 85 | 81.04 | 9.16 | 9.81 |
| CK22 | 44432270 | 99.28 | 0.7 | 8.55E-05 | 0.02 | 91.63 | 84 | 80.06 | 10.15 | 9.79 |
| CK23 | 45263166 | 99.06 | 0.91 | 0.02 | 0 | 92.17 | 82 | 81.5 | 9.04 | 9.46 |
| T11 | 452252<br>88 | 99.07 | 0.91 | 0.02 | 0 | 92.41 | 82 | 81.03 | 9.22 | 9.75 |
| T12 | 476942<br>60 | 98.96 | 1.02 | 0.02 | 0 | 92.13 | 86 | 82.08 | 8.45 | 9.47 |
| T13 | 477606<br>84 | 99.29 | 0.68 | 0 | 0.03 | 89.68 | 81 | 79.47 | 10.21 | 10.33 |
| T21 | 452518<br>76 | 99.11 | 0.87 | 0.02 | 0 | 92.46 | 76 | 81.01 | 9.35 | 9.64 |
| T22 | 474777<br>02 | 98.99 | 0.98 | 0.02 | 0 | 92.32 | 83 | 81 | 9.15 | 9.85 |
| T23 | 458495<br>30 | 99.21 | 0.77 | 0.02 | 0 | 92.38 | 80 | 81.66 | 8.86 | 9.48 |

C: control groups without GA treatment
T: GA treatment groups
Ns reads rate means as the N is too high, the number of sequences that have been removed accounts for the proportion of the original sequence number.

### Identification and classification of DEGs at different stages

In the two key developmental stages of European sweet cherry, changes in gene expression were determined by comparing C1 versus T1 and C2 versus T2. In these two periods, 765 and 186 differentially expressed genes were obtained. In the first period, 685 genes were upregulated, and 76 genes were downregulated. In the second, 141 upregulated and 45 downregulated genes were identified by RNA-seq (Figure 3A). The number of upregulated genes is significantly higher than the number of downregulated genes in the two periods of European sweet cherry development. Venn diagram analysis indicates that only 32 genes showed a pattern of upregulated expression in both key periods (Figure 3B). In addition, at every time point, the specific upregulated genes can be detected.

**Fig. 3.**
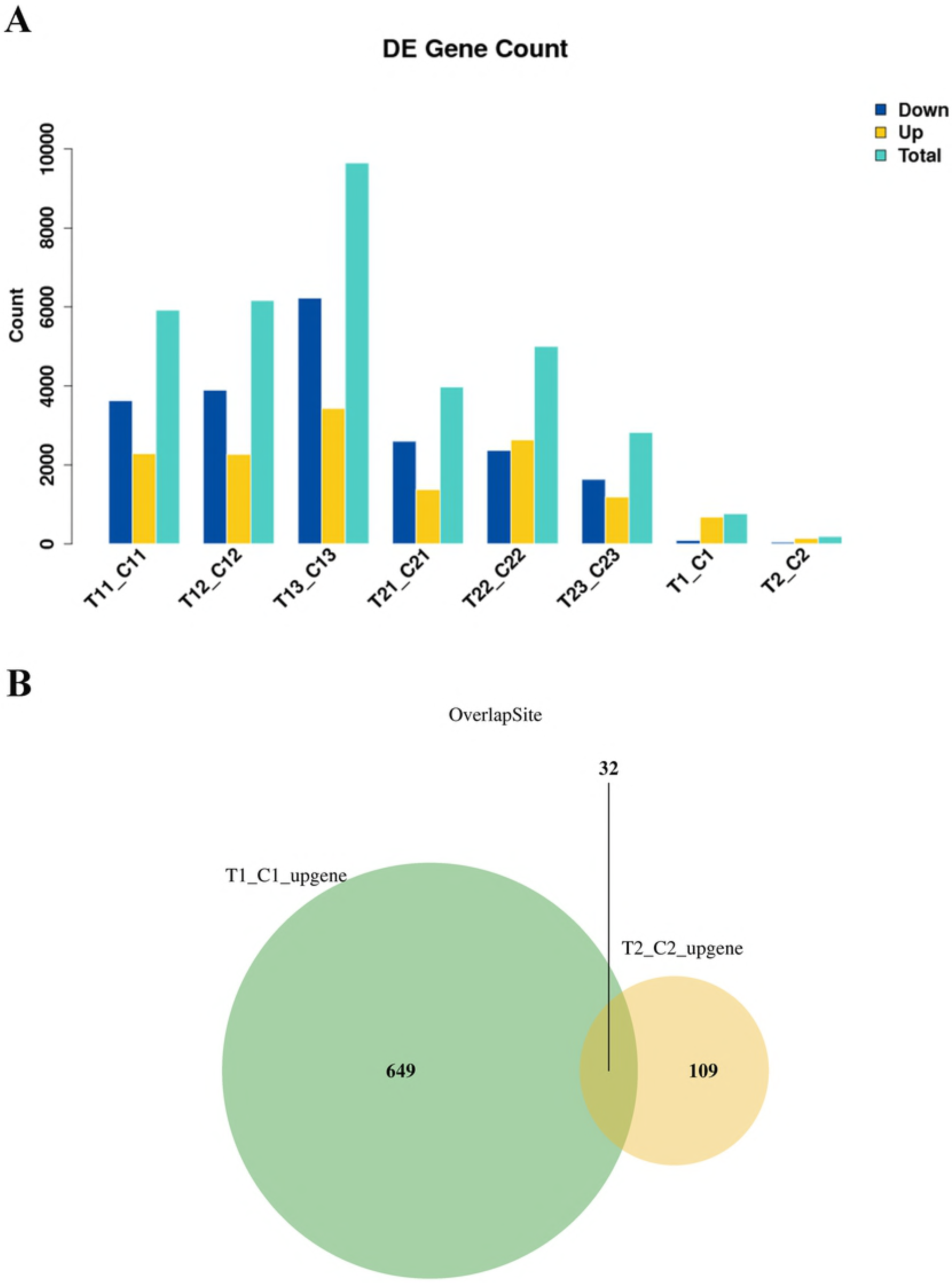
Comparison of differentially expressed genes (DEGs, *P* value≤0.05) at two key sweet cherry developmental stages. Numbers of DEGs between control and GA3 treatment samples at two dvelopmental stages (A); Venn diagram representing the relationship between up DEGs identified in two sweet cherry development stages (B).

### Changes in the expression patterns of gibberellin metabolism and signal transduction genes

After the treatment with gibberellin, the fruit rate, fruit size and rate of parthenocarpy significantly changed. Gibberellin 2-beta-dioxygenase (*GA2ox*) is a key enzyme of GA catabolism [26]. During the anthesis period, the gene expression level of *GA2ox* changed significantly, and it was also detected during the hard-core period. Pav_sc0000095.1_g1110.1.mk was significantly upregulated after 7 days (Table S2), and its annotation was *PavGA2ox*, which was used to generate GA51, GA29, GA34 and GA8 in GA9, GA20, GA4 and GA1 respectively [27]. In *Arabidopsis thaliana*, the *Scarecrow-like* (*SCL*) and *DELLA* proteins are antagonists, and they promote the transduction of the gibberellin signal [28]. The transcriptome sequencing detected some genes at the anthesis stage after GA treatment, such as *pavscl1, pavscl3* and *GA2ox*. A GA receptor gene, *GID1B* (Pav_sc0000848.1_g080.1.mk), was downregulated after GA3 treatment in the hard nucleus phase, and *GA2ox* and *DELLA* were upregulated. The results of this study showed that exogenous GA3 could be associated with parthenogenesis.

In the GA signal transduction pathway, active GA regulates the hydrolysis and inactivation of *DELLA* protein by stimulating the GID1-DELLA-GID2 complex, thus regulating downstream gene expression [29]. In this study, we found that the gene encoding *GID1* was downregulated after GA treatment, the gene expression of the encoded *DELLA* protein was upregulated, and the gene encoding *GID2* was not detected. Among them, Pav_sc0000848.1_g080.1.mk was further annotated as *PavGID1B* and only noted in the hard-core period. During the hard-core phase, the *DELLA* protein Pav_sc0000464.1_g350.1.mk significantly increased. The above results indicated that after the treatment of ‘Meizao’ sweet cherries with exogenous GA3, no significant expression of GID1-DELLA-GID2-related genes was detected at the full anthesis stage, and we speculated that the GID1-DELLA-GID2-related genes had responded very quickly and were correlated with seed development.

### Changes in expression patterns of cytoskeleton and cell wall modification genes

After GA3 treatment, significant changes took place in the size and shape of the ‘Meizao’ cherries. After 7 days, the treated fruit was significantly swollen (Figure 2). Upregulation of the plant hormone synthesis and decomposition, control and signal transduction genes and downregulation of genes involved in the cytoskeleton, cell wall modifications and other related biological processes synergistically caused irreversible fruit expansion after processing (Table S3). The expression of the genes encoding actin and tubulin changed significantly after GA3 treatment. After 7 days of anthesis, only one gene encoding actin, *EDH1* (Pav_sc0001084.1_g180.1.mk), was downregulated. The genes encoding tubulin were not detected during the hard-core period. The different genes involved in cell division, including a cyclin-dependent kinase and five histones, were downregulated at the anthesis stage after GA treatment.

The expression of cell wall modification genes was significantly changed after treatment. Studies have shown that the expansion proteins can catalyze the extension of cell walls [30]. In our study, 26 cell wall-associated DEGs were detected at the anthesis stage after GA treatment, and 5 cell wall-associated DEGs were detected during the hard-core period, including those encoding expansin, xyloglucan glycosyltransferase and glucanase, and among them, most of these genes (27/31) were upregulated. It is clear that the changes in the genes associated with cell division and cell wall metabolism are the basis of fruit development.

### DEG analysis of transcription factors

By analyzing all the DEGs at the two stages, we obtained 403 transcription factors, further divided into 46 and 30 gene families (Figure 4A). In the first period, the families of transcription factors with the most DEGs are the bHLH and B3 families (7.94%), followed by the NAC family (5.96%); the WRKY family (4.96%); the MYB-related family (4.71%); the C3H, C2H2, ERF and bZIP families (4.47%); the FAR1 family (4.22%); the MYB family (3.97%); the HSF family (3.23%); the TALE family (2.98%); the ARF family (2.73%); and the M-type, HB-other, and TCP families (2.23%). These 17 families of transcription factors contained a higher than average number of genes. In the second stage, B3 was the family of transcription factors with the most DEGs (12.9%), followed by the NAC, bHLH and HD-ZIP families (7.52%); the C3H family (6.45%); the C2H2 family (5.38%); and the MYB-related, GRAS and MYB families (4.3%) (Figure 4B). The abovementioned families of transcription factors contained a higher than average number of DEGs. By analyzing the expression patterns of the DEGs, it was found that the transcription factors of most of the DEGs showed an upregulated trend compared with the control at the two key stages of gibberellin treatment. While some of the DEGs belonging to the Nin-like, HSF, ARF, B3, HB-other, MYB, Bzip, GeBP, C3H and NF-YC transcription factor families were downregulated after the first treatment, the NAC and Trihelix families were downregulated after the second treatment. As previous studies have shown, many transcription factors play an important role in the development of a wide variety of plant seeds [31]. For example, the B3 superfamily plays a key role in seed development [32–34]. Moreover, all the NAC genes were reported to be involved in the synthesis of cell secondary metabolites [35]. In addition, transcription factors (TFs) such as GRAS and HB participate in the signal transduction of ABA and GA [36] (Figure 5,6). In our study, it was found that the transcriptional factors related to seed development mentioned in the above examples were differentially expressed between seeded and parthenogenetic progeny, indicating that they are associated with parthenogenetic phenotypes.

**Fig. 4.**
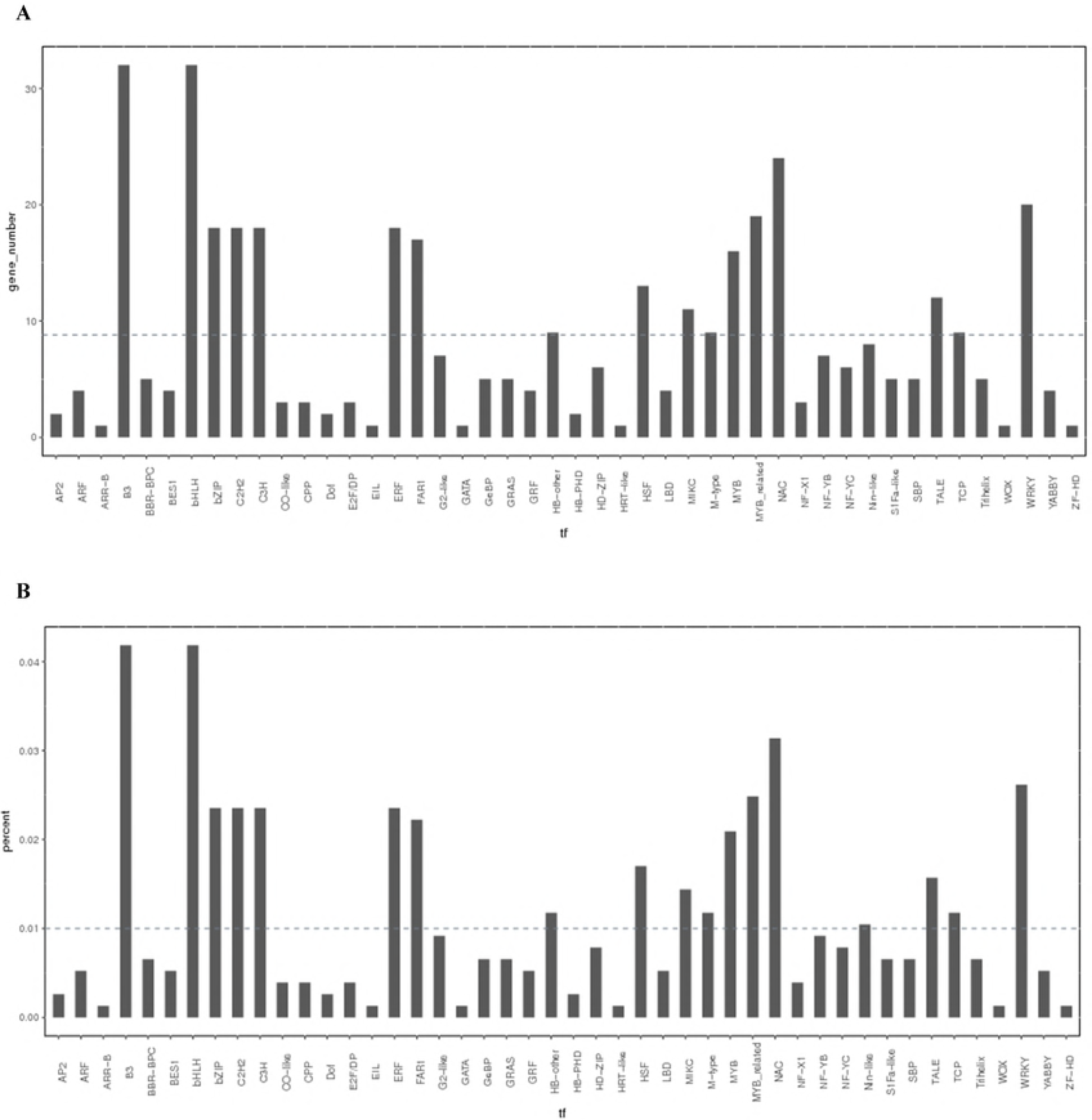
The number of DEGs in the family of different transcription factors, dashed line indicates the average number of DEGs in different families (A); The proportion of the different transcription factor family to the total transcription factor familys (B);

**Fig. 5.**
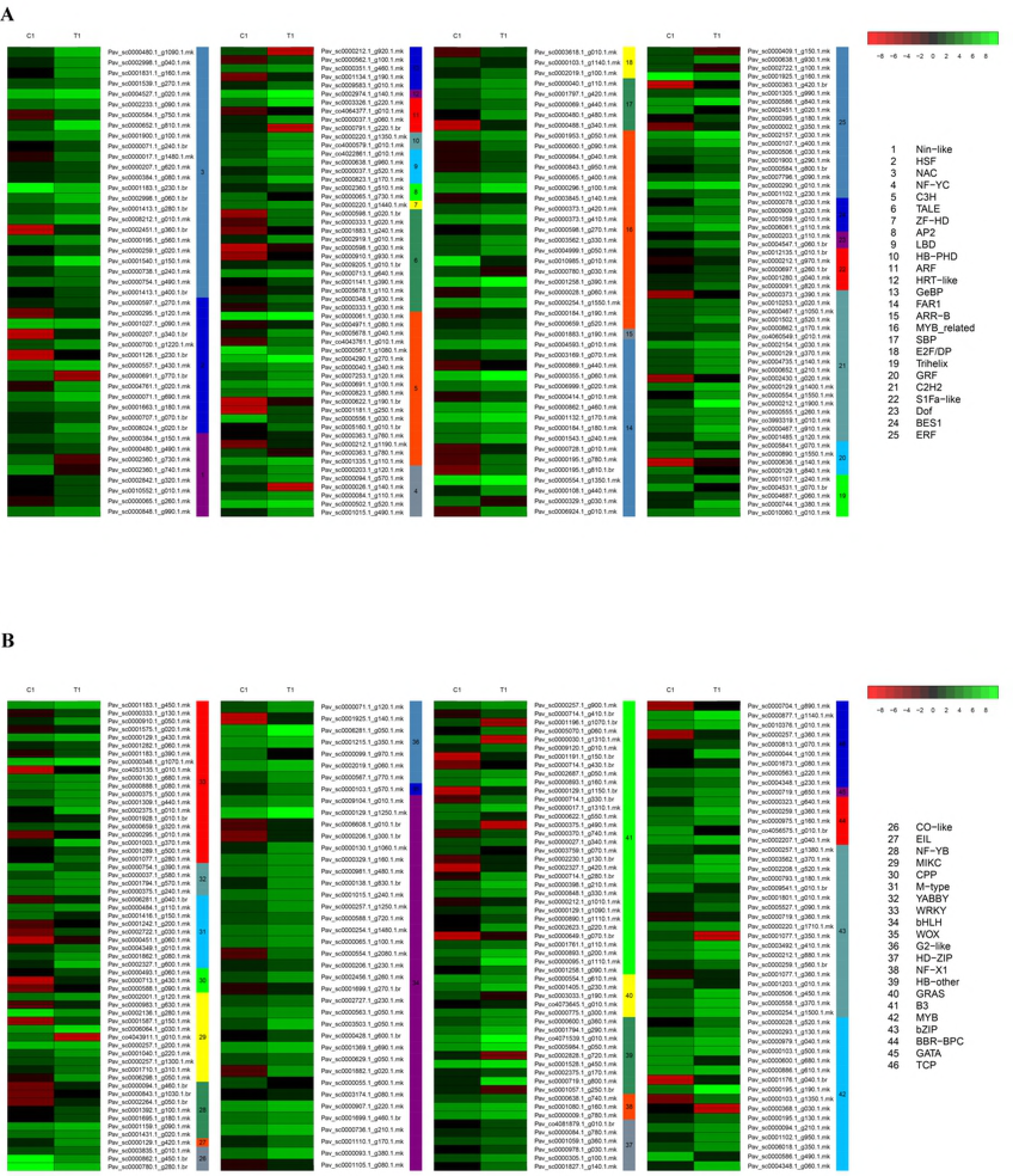
Expression analysis of genes from different transcription factor familys (T1/C1).

**Fig. 6.**
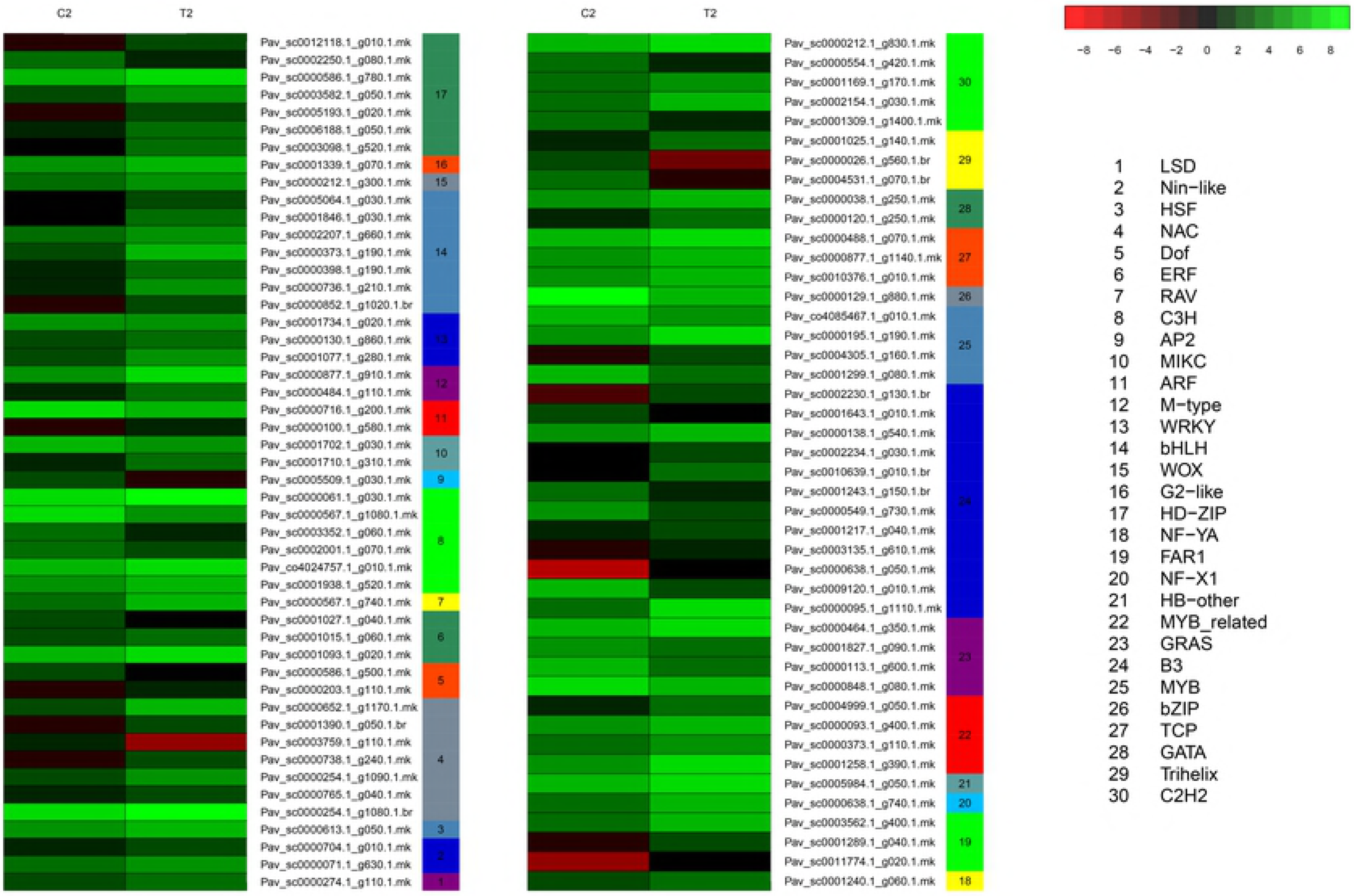
Expression analysis of genes from different transcription factor familys (T2/C2).

### Analysis of DEG pathway enrichment to reveal the possible pathways influencing seed development

To further understand the difference in the metabolic pathways of ovule development between seeded and parthenogenetic sweet cherry, we analyzed the metabolic pathways with differential gene enrichment in the two stages (Figure 7). The results suggested that a total of two metabolic pathways were significantly enriched at the first stage of cherry development. However, no pathways were significantly enriched at the second developmental stage. Two metabolic pathways, phenylalanine metabolism and phenylpropanoid biosynthesis, were significantly enriched at the first stage of cherry development, and most of the DEGs were upregulated in these two pathways. Collectively, these results suggest that parthenogenetic fruit set may be related to the phenylalanine metabolism and phenylpropanoid biosynthesis pathways.

**Fig. 7.**
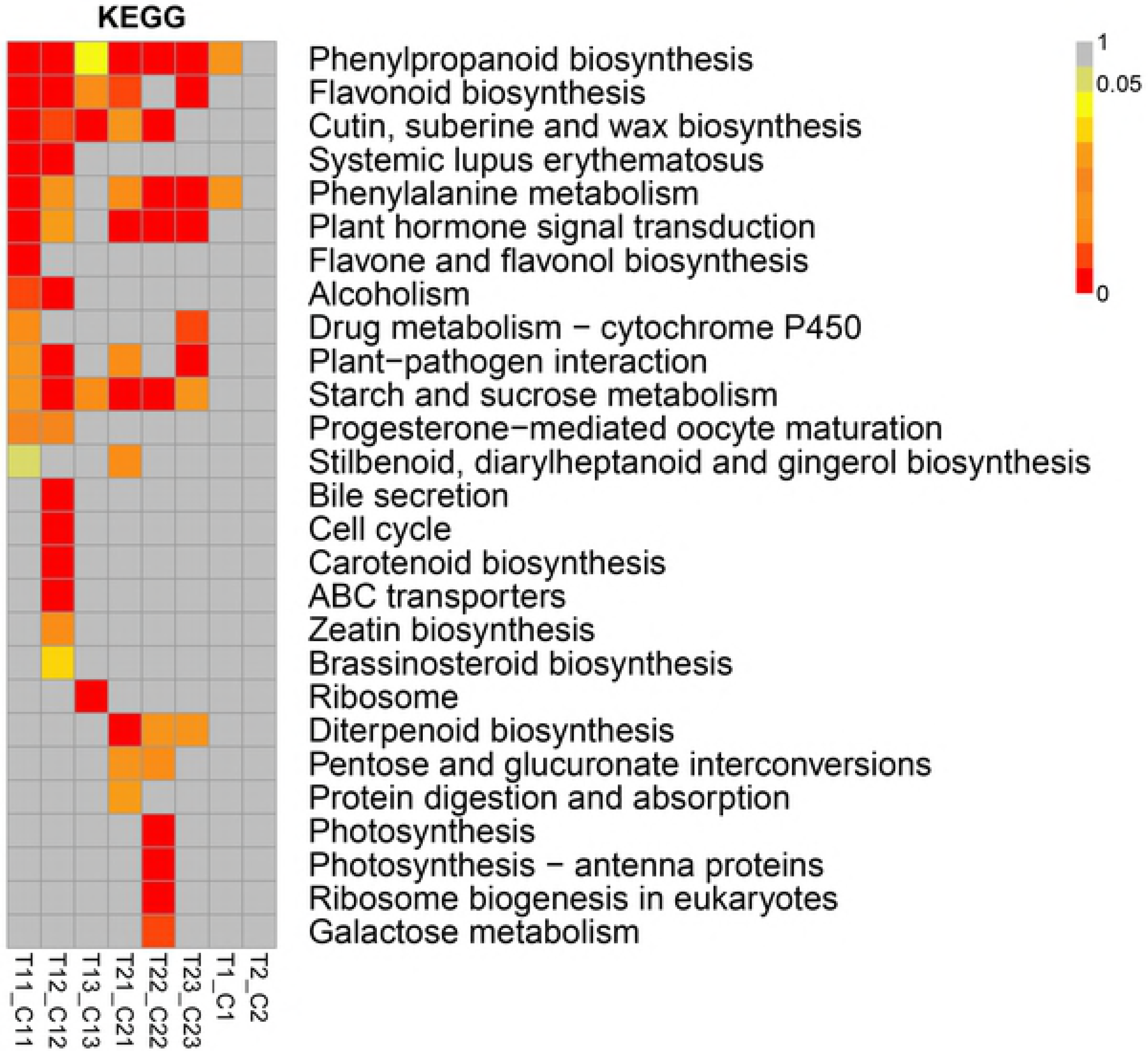
Analysis of DEGs metabolic pathway enrichment at two different developmental stages of sweet cherry.

### Functional analysis of GOs of differential genes indicates the factors that may lead to parthenogenetic fruit set

To explore the key biological processes and functional classification related to cherry parthenogenetic fruit set, the DEGs were further analyzed for biological function GOs in the two critical stages of cherry development (Figure 8). The biological functional classification of GOs mainly involves the following three categories: biological process, molecular function and cellular component. Most of the DEGs were upregulated, and the ones detected at the first stage have high representation in the cellular process, metabolic process, single-organism process, response to stimulus, biological regulation, and cellular component organization of biogenesis groups in the biological process category. In general, most of the differences in gene-rich GOs are mainly related to plant growth and development, response to external stimuli and signal transduction, glucose and lipid metabolism, epigenetic regulation, etc.; the abovementioned groups and parthenogenetic fruit set are closely related. In addition, in the anthesis stage after GA treatment, the status of differential gene enrichment is generally the same.

**Fig. 8.**
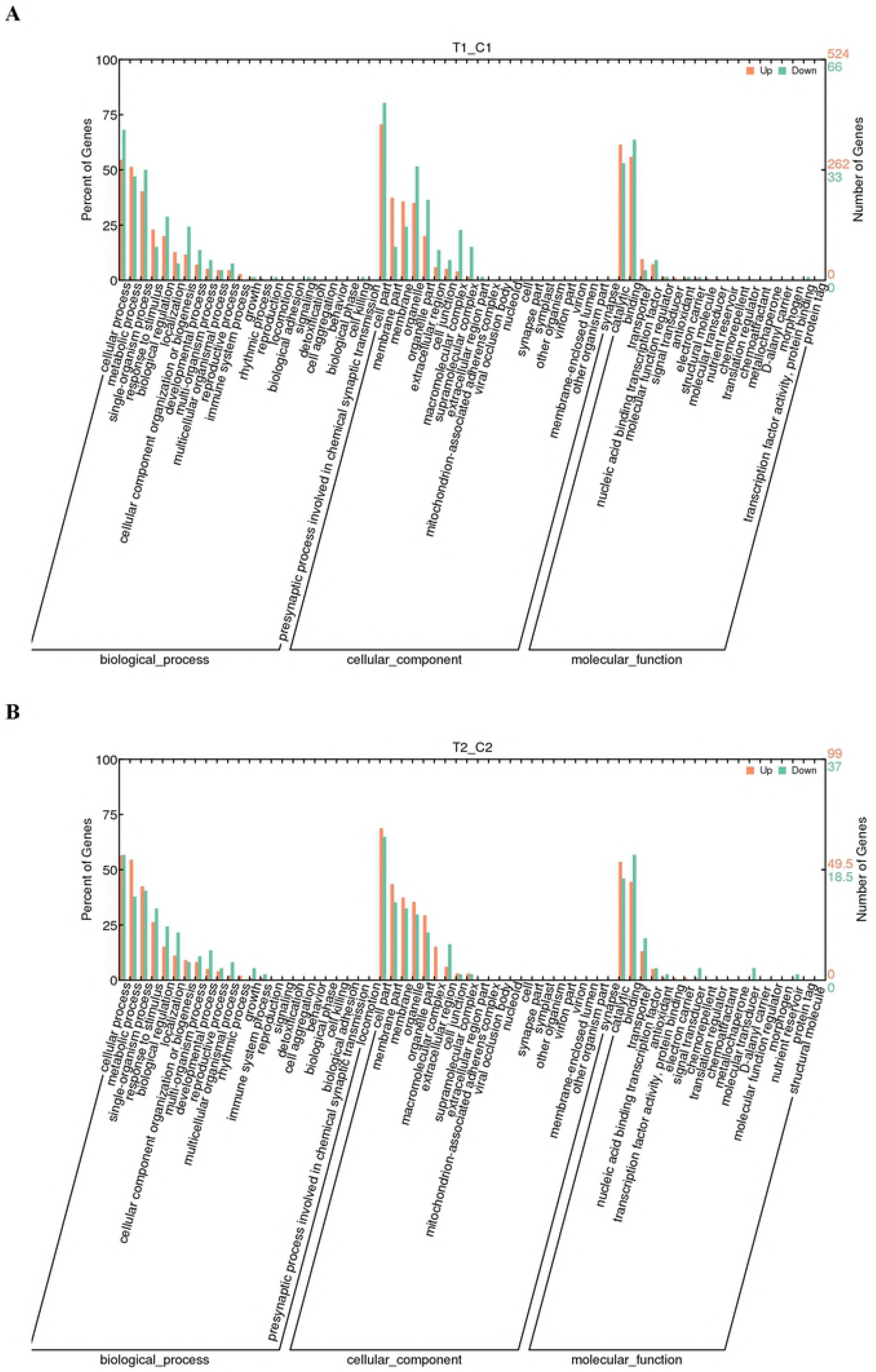
Gene ontology (GO) (corrected *P*-value≤0.05) analysis of control and GA3 treatment.biological component; cellular component; molecular function. The numbers on the columnar represent the percentage of the difference genes annotated to each GO terms in total DEGs. T1/C1 (A), T2/C2 (B).

Based on the classification of GO-based molecular functions, most of the DEGs are enriched in catalytic, binding, transporter and nucleic acid-binding transcription factors. These enriched GOs are closely related to the regulation of biological processes and the anabolism of functional substances. In each subcategory, more genes were upregulated than downregulated in the first stage of sweet cherry development.

Analyses were performed on the GO classification of cellular components, and most DEGs are enriched in the following groups: intrinsic component of membrane, integral component of membrane, membrane and membrane part. Similar to the GO enrichment of molecular function and biological process, in the GO enrichment of the cell group, most of the DEGs were upregulated in the anthesis stage after GA treatment compared with the control.

To further explore the differences in the GO functions enriched in the DEGs during the two critical stages of sweet cherry development, the analysis focused on the GO enrichment of the biological process category. During the two critical stages of sweet cherry development, a total of 709/24 upregulated and 55/0 downregulated genes were found to be significantly enriched compared to the control. The GO items enriched in these two developmental stages were further summarized. In the anthesis stage, it was found that the GO items with upregulated DEGs were mainly associated with cell wall development. This can be further divided into the groups of cell wall synthesis and cytokinesis, carbohydrate and polysaccharide biosynthesis, secondary metabolism, lignin and phenylpropanoid anabolism. However, the downregulation of GO items related to cell wall development was associated with DEGs significantly enriched in the cell cycle process during the anthesis stage. It was found that the GO items with upregulated DEGs were mainly associated with photosynthesis and sucrose biosynthesis, and no downregulated genes were enriched in the second stage.

### Transcriptional dynamics analysis of DEGs

After GA3 treatment, in order to analyze the genes associated with vertical diameter and parthenocarpy in the different developmental stages of CK sweet cherries one at a time, this study performed individual comparisons of the vertical diameter and parthenocarpy of sweet cherries in the CK treatment (Figure 9). The different genes (15167 in total) were subjected to k-cluster analysis of temporal and spatial expression patterns. Through the k-cluster analysis, all the differentially expressed genes with similar expression patterns during the process of cherry development were grouped into one group, and the results are shown in figure 7. Through the k-cluster analysis, a total of six coexpressed clusters were identified during the process of the development of sweet cherries in the CK and the GA3 treatment, of which 4 were consistent in expression pattern. The DEGs were consistently upregulated in the CK for the parthenogenetic rate of sweet cherry at the early stage of development in cluster 1, cluster 2, cluster 4, and cluster 6, and the only difference was the degree of upregulation. These four clusters showed lower expression in the anthesis period after GA treatment. Cluster 5 showed different expression patterns, and the transcriptional level of the DEGs showed continuous downregulation. After GA treatment, the DEG patterns of cluster 5 and clusters 1, 2, 4, and 6 were reversed and correlated through time. The transcription level was increased or decreased during the critical period of sweet cherry development (anthesis period), and the differential genes between the five clusters had completely opposite spatiotemporal expression patterns during the development of sweet cherries in the CK and the GA3 treatment.

**Fig. 9.**
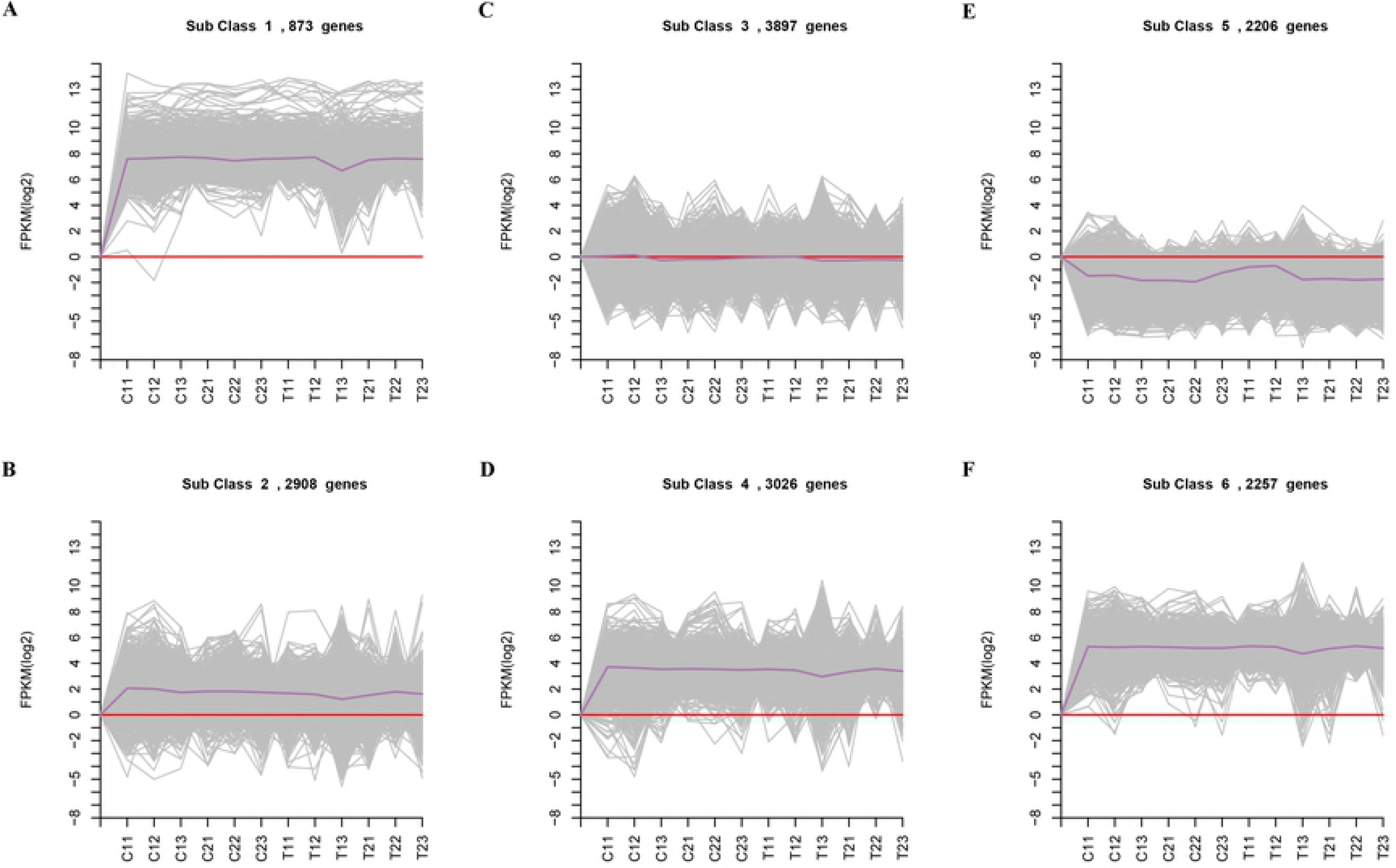
Expression Pattern Clustering of DEGs at two key Sweet Cherry Development stages with Control and GA3 treatment. Clustering using k–means method. The Numbers behind each cluster name represent the number of different genes in the cluster. The X–axis shows the period 2 versus period 1 of the control and treatment. The Y–axis shows the log2–value of RPKM during the development of sweet cherry.

Based on the clustering results, 5 of the transcription factors were analyzed, and it was found that the highest number of transcription factors in cluster 5 belongs to the AP2 and MYB families, followed by the MADS family. In cluster 5, the different genes are mainly enriched in the GO items of seed and embryo development, cell differentiation, cell wall formation, defense response, hormone regulation, transcriptional regulation, etc.

### RNA-Seq validation of DEGs by Quantitative real-time PCR (qRT-PCR)

To validate the reliability of the transcriptome data, we detected the expression of 30 DEGs related to seed development based on gene annotation and the existing literature with homology analysis using qRT-PCR in samples from the “treatment” and the control (Figure 10). We further carried out a correlation analysis of gene expression fold changes under the two methods. As shown in Figure 8, the results were highly consistent and thus verified the reliability of the transcriptome sequencing data. These 32 genes are mainly involved in fruit setting, such as *PavYUCCA*, *PavYUCCA10, PavACS, PavACO* and *PavCYP707A3*; cell division, such as *PavH4, PavCDC2D, PavH3.2, PavCDC45*, etc.; and parthenocarpy, such as *PavSAUR32*, *PavSCL1*, *PavCYCPA3*, *PavDELLA*, etc. The sample for qRT-PCR was a mixed sample of the sweet cherries obtained during the two critical periods of sweet cherry development and used for the transcriptome sequencing. The qRT-PCR results for the 30 differentially expressed genes in the sweet cherry samples used for transcriptome sequencing are shown in figure 8 and are consistent with the results from transcriptome sequencing.

**Fig. 10.**
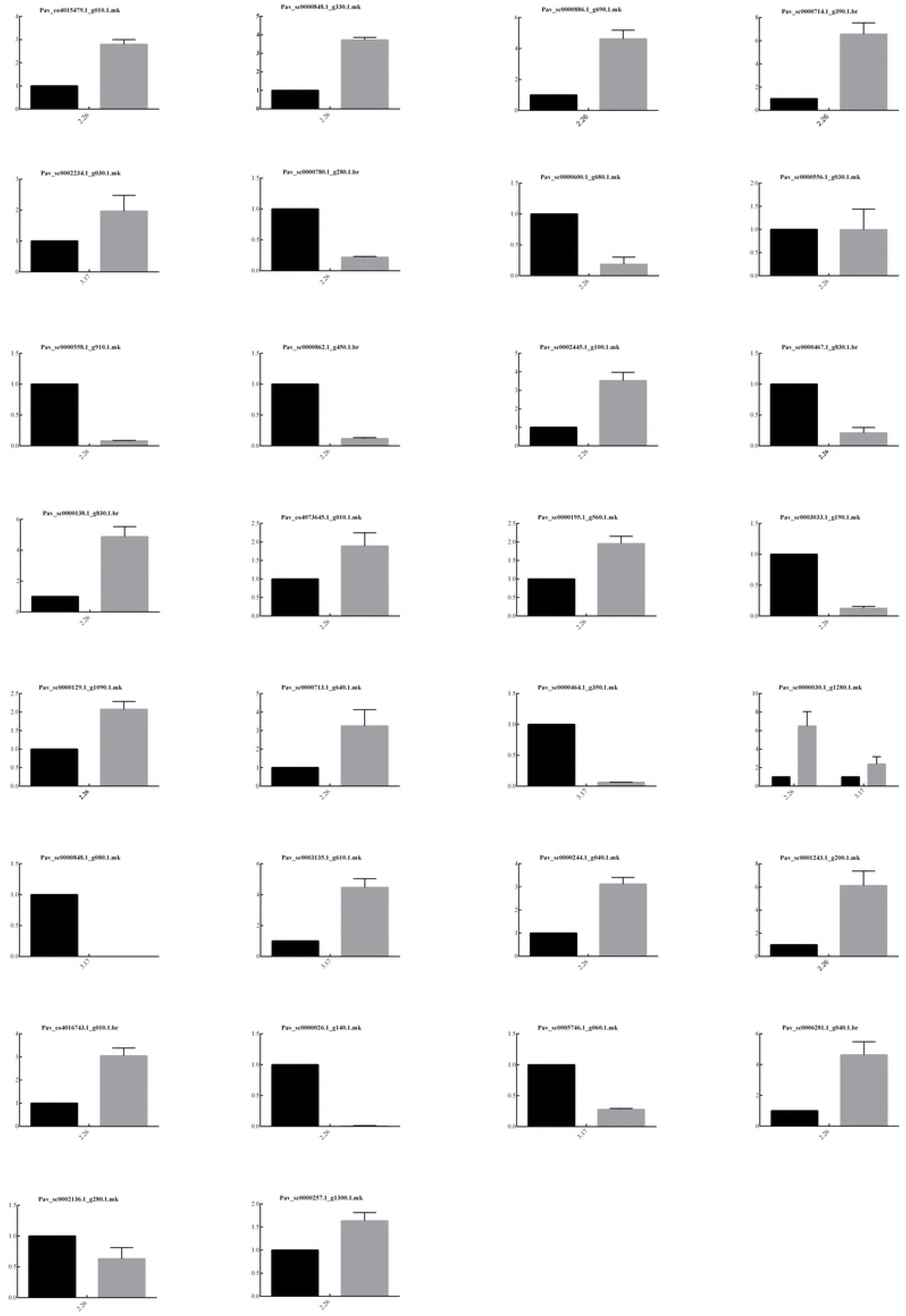
Transcriptome results were verified by qRT-PCR.

## Discussion

Transcriptome analysis was performed during the growth and development of ‘Meizao’ cherry in response to GA3 during two critical periods of sweet cherry development. In total, 765 and 186 differentially expressed genes were identified, and they were then compared to determine their biological function based on GOs and the metabolic pathways that were enriched. In this study, GA3 treatment was carried out at two key stages of sweet cherry development, a one-to-one comparison of parthenocarpic fruit development after GA3 treatment was conducted, and the differences during the fruit developmental period were compared, resulting in a relatively comprehensive follow-up analysis conducive to the identification of important genes linked to sweet cherry fruit production that participate in functional pathways and to the exploration of the potential mechanisms regulating and controlling the production of sweet cherry fruit.

### The effect of gibberellin gene expression on the development of Parthenogenetic sweet cherry

In agriculture, the production of parthenocarpic seedless grapes often involves treating flower buds with gibberellin before flowering [37]. After the treatment with exogenous GA3, a GA2ox gene was detected to promote GA inactivation during the first period, and two *GA2ox* genes were detected to promote GA inactivation during the second period. Therefore, the analysis of hormone-related gene expression during the development of sweet cherry in two stages can help to further analyze the influence of gene expression differences on the parthenocarpy of sweet cherry and then explore the mechanism of parthenocarpy formation. Previous studies have shown that *SAUR* can promote the expansion of *Arabidopsis* cells [38,39], and *SAUR* was upregulated in the first stage in the IAA signal transduction pathway, which is consistent with the results shown in Figure. 2. Expression of DEGs induced parthenocarpy by GA3 were showen in Table S2. The differentially expressed genes associated with auxin synthesis and the auxin inhibitor IAA also showed patterns of upregulated expression at the first stage. It can be presumed that these auxin-related differential gene expression patterns may be a manifestation of auxin balance feedback adjustment. The *GH3* gene has the function of maintaining the dynamic balance of auxin in plants [40]. In the hard-core stage of sweet cherry, the expression of the *GH3* gene and the auxin inhibitor *AUX/IAA* was downregulated, which further explained the balance feedback regulation of auxin. Different hormones can play a cooperative role through signal exchange, and the hormone that acts synergistically with auxin is BR. Studies have shown that auxin promotes plant growth by synthesizing BR [41].

In the response pathway of BR, the upregulated expression of the receptor gene *BRI1* indicates a connected effect on the GA-induced BR response. In this study, the genes related to BR synthesis showed upregulated expression in two critical stages of sweet cherry development, which is consistent with the synergistic effect of auxin and brassinolide. The study of Fu showed that external application of BR can stimulate cucumber’s cell division and lead to parthenocarpy [42]. It is speculated that the expression of genes related to BR synthesis and signal transduction in sweet cherry during seed development after gibberellin treatment is upregulated relative to that in untreated sweet cherry. It may also promote the parthenogenesis of sweet cherry through similar mechanisms.

It is reported that the high level of ethylene response can cause the parthenogenesis of tomato [43]. The results showed that in the first period, the expression of genes related to ethylene synthesis and signal transduction was upregulated after GA3 treatment. Studies have shown that abscisic acid participates in the development of the seed coat in *Arabidopsis* and regulates the metabolism of cell wall by inhibiting the expression of PG, PE and XET in the cell wall [44,45]. In the first critical period of development of sweet cherry, the expression of a gene related to seed growth was detected by analyzing the expression pattern of seed development-related genes, which was consistent with the upregulated expression of the abscisic acid receptor and the gene expression related to transmembrane transport at the two critical periods of sweet cherry development. However, no genes associated with seed coat development, which may only be expressed a short time after treatment, were detected in the second period. In summary, the genes associated with GA, auxin and BR were significantly upregulated after GA3 treatment, which may be associated with parthenogenesis, and its specific mechanism needs further study.

### Gibberellin promotes cell wall relaxation and induces fruit enlargement

The cell wall is composed of cellulose microfilaments embedded in the hemicellulose stromal layer through noncovalent bonding [46]. In the process of fruit enlargement, the expansion proteins promote noncovalent dissociation, while glucanase and xylanase can loosen the hemicellulose matrix [47]. Expression of DEGs associated with cell division after GA3 treatment were showen in Table S3. In our results, glucanase gene ncRNA was significantly upregulated in the first stage, and the endoglucanase gene *ENDO17* was upregulated in the second stage after GA3 treatment. The two expansion genes *EXPA2* and *EXPB1* were significantly upregulated in the anthesis period, but only one gene, *EXPA10*, was detected in the second period, and its expression was downregulated. The glucanase, xylanase and expansion genes were significantly upregulated in the first period, and one expansion gene was downregulated in the second period after GA3 treatment, suggesting that these genes play an important role in the process of sweet cherry enlargement. Previous studies have found that the *XET* gene was significantly upregulated during the first rapid growth stage of the grape and then decreased over time [48,49], which is not consistent with the experimental results. The main reason may be that the expression patterns of different species are different. The growth and development period of sweet cherry is short. After GA3 treatment, the rapid growth period of sweet cherry is relatively long, and the glucanase, xylanase and expansion genes are highly expressed. It is known that cell wall modification plays a key role in exogenous GA3-induced fruit enlargement. Compared with the genes associated with cell wall relaxation, the cytoskeleton modification genes also play an important role in the process of GA3 inducing fruit enlargement. The expression of 3 actin and 10 tubulin genes was upregulated, according to the analysis of the protein groups in the second and third period of grape development.

### Gibberellin-Induced fruit setting in sweet cherry

During plant growth and development, pollination is a key step in the transition from flowers to fruits. The egg cells are induced into seeds by pollen fertilization, and the zygotes stimulate the ovary to form fruit. In the current study, three genes may play an important role in the fruit setting of sweet cherry. First, the ethylene synthesis pathway enzymes 1-aminocyclopropane-1-carboxylate oxidase (*ACO*) and 1-aminocyclopropane-1-carboxylate synthase (*ACS*) were upregulated after GA treatment. The effects of ethylene include promoting fruit ripening, and ethephon is used as a thinning agent for many fruit trees [50–52]. The high expression of these two genes may induce the synthesis of ethylene and subsequently induce the loss of fruit. Second, it is reported that the *YUCCA* gene plays an important role in the biosynthesis of auxin and the development of plants [53]. In this study, expression of DEGs associated with fruit setting after GA3 treatment were showen in Table S4. The indole-3-pyruvate monooxygenase *YUCCA10* was significantly upregulated after GA treatment. The *YUCCA* gene plays an important role in vegetative growth and reproductive development of plants and regulates the formation of the floral meristem and the development of lateral organs [54]. Some studies have reported that the content of auxin in the ovary increased after GA treatment [55–56]. GA may play a role in the accumulation of auxin in the ovary and in turn act as a signal to affect fruit set. Third, the ABA 8’-hydroxylase gene was upregulated after GA treatment in the second stage. This gene is thought to be the main reason for the decrease in ABA [57]. High ABA content causes the fruit to fall off. After GA treatment, the expression of the abscisic acid 8′-hydroxylase gene was upregulated, which was beneficial to the fruit. The results are consistent with the results shown in figure 2, indicating that this period is a critical period for fruit setting. These results indicate that auxin, ethylene and abscisic acid may regulate fruit setting in sweet cherry after GA treatment.

## Conclusion

To the best of our knowledge, this study is the first to provide a dynamic analysis of the differences in the differentially expressed genes related to parthenogenesis, fruit enlargement and fruit setting of sweet cherry fruit caused by GA3 treatment. We selected two critical periods during the development of sweet cherries, and the genes of many metabolic processes changed significantly after a week of GA treatment. In the GA signaling pathway, these included hormone-related genes, transcription factors and cell division genes, and the further analysis of these genes will provide us with a new way to study the fruit setting of sweet cherry. At the same time, our research results also provide a relatively complete molecular platform for future research on mechanisms of parthenocarpy, fruit enlargement and fruit setting in sweet cherry.

## Supplementary data

Table S1 The primers used in this article.

Table S2 Expression of DEGs induced parthenocarpy by GA3

Table S3 Expression of DEGs associated with cell division after GA3 treatment.

Table S4 Expression of DEGs associated with fruit setting after GA3 treatment.

## Acknowledgement

Thanks to DSG and WX for helping me designed the research. WX, ZJZ and WLS helped to analyzed the physiology data. MYS helped to designed the qRT-PCR primes. BBW, SXL and CM revised the intellectual content of this manuscript.

